# Altered Huntingtin-Chromatin Interactions Predict Transcriptional and Epigenetic Changes in Huntington’s Disease Mouse Models

**DOI:** 10.1101/2020.06.04.132571

**Authors:** Jocelynn R Pearl, Amol C Shetty, Jeffrey P Cantle, Dani E Bergey, Robert M. Bragg, Sydney R. Coffey, Holly B. Kordasiewicz, Leroy E Hood, Nathan D Price, Seth A Ament, Jeffrey B Carroll

## Abstract

Progressive striatal gene expression changes and epigenetic alterations are a prominent feature of Huntington’s disease (HD), but the mechanistic basis remains poorly understood. Using chromatin immunoprecipitation and sequencing (ChIP-seq), we show that the huntingtin protein (HTT) reproducibly occupies specific locations in the mouse genome. Moreover, many genomic loci were differentially occupied by HTT in striatal tissue from a homozygous knock-in mouse model of HD (B6.*Htt^Q111/Q111^*) when compared to wildtype controls. Huntingtin ChIP-seq peaks were enriched in the coding regions of cell identity genes important for striatal function, with many of these genes found to have reduced expression in the striata of HD patients and mouse models, as well as reduced HTT occupancy in *Htt^Q111/Q111^*mice compared to controls. Conversely, HTT ChIP-seq peaks were depleted near genes that are up-regulated in HD. ChIP-seq of bulk striatal histone modifications, generated in parallel, revealed genotype-specific colocalization of HTT with active chromatin marks (H3K4me3 and H3K27ac), and with enhancer of zeste homolog 2 (EZH2), a key enzymatic component of the PRC2 complex. Near genes that are differentially regulated in HD, greater HTT occupancy in *Htt^Q111/Q111^* vs. wildtype mice was associated with increased EZH2 binding, increased histone H3 lysine 4 (H3K4me3), and decreased histone H3 lysine 27 (H3K27me3). Our study suggests that huntingtin-chromatin interactions may play a direct role in organizing chromatin and promoting cell type-specific gene expression, with loss of HTT occupancy predicting decreased gene expression in HD.

## Introduction

Huntington’s disease (HD) is a fatal dominant neurodegenerative disease caused by expansion of a glutamine-coding (polyQ) CAG tract near the 5’ end of the *Huntingtin* (*HTT*) gene[1]. Clinical symptoms include deficits in motor control and cognition, as well as psychiatric symptoms. HD progression is linked to the selective death of spiny projection neurons (SPNs) in the striatum[2]. Transcriptional[3] and epigenomic[4–7] dysregulation are among the earliest phenotypes in cells and tissues expressing mutant HTT protein (mHTT) and are highly reproducible in brain tissue from humans with HD[8–10]. However, the molecular mechanisms by which mHTT mediates these transcriptional changes are poorly understood.

A straightforward hypothesis is that HTT directly contributes to transcriptional dysregulation in HD through interactions with chromatin-bound DNA. In the cell, HTT protein dynamically shuttles between the cytoplasm and nucleus[11]. Mis-localization of mHTT to the nucleus is an early phenotype in HD animal models, roughly coincident with the onset of transcriptional changes, and is a critical driver of mHTT-mediated neuronal death, both *in vitro* and in mouse models[11–13]. To date, specific interactions of HTT with chromatin remain obscure, but they are supported by several lines of indirect evidence: (i) chromatin immunoprecipitation indicates that HTT affiliates with chromatin DNA[14]; (ii) the HTT protein contains a series of HEAT domains, which are capable of serving as DNA binding domains[15]; and (iii) HTT forms direct protein-protein interactions with a variety of transcriptional regulatory proteins in the nucleus, including transcription factors (TFs) and chromatin remodeling factors[16]. Perhaps the best understood of such interactions involve chromatin remodeling complexes that mediate gene repression and heterochromatin formation. HTT binds in a polyQ-length sensitive manner to polycomb repressive complex 2 (PRC2), the chromatin remodeling complex responsible for trimethylation of lysine 27 on the histone 3 tail (H3K27me3), a repressive mark associated with bi-valent, or poised chromatin regions that are critical to developmental fate commitment[17]. Ablation of PRC2 in SPNs reproduces several of the cellular phenotypes in HD, including aberrant de-repression of transcripts that code for developmentally regulated transcription factors, loss of SPN identity gene expression, and prolonged cell death[18]. We[19] and others[4,10] have observed that a common transcriptional feature of HD is a reduction of cell-type appropriate gene expression, providing additional impetus to understand changes in PRC2 function in HD, given its key role in the regulation of cell fate[20].

To investigate the hypothesis that HTT occupies specific locations on chromatin DNA, we performed chromatin immunoprecipitation and deep sequencing (ChIP-seq) to map HTT genomic occupancy in striatal tissue from the *Htt^Q111/Q111^*knock-in mouse model of the HD mutation and in wildtype *Htt^+/+^*controls. We analyzed these data together with publicly available RNA-seq and with newly generated ChIP-seq of the histone modifications H3K27me3, H3K9me3 and H3K4me3, and the PRC2 histone methyltransferase effector subunit EZH2, generated in parallel from the striatum of age-matched of *Htt^Q111/+^* mice. We describe thousands of reproducible HTT ChIP-seq peaks, both in *Htt^+/+^*and *Htt^Q111/Q111^* mice, and observe robust genotype-specific patterns of occupancy, which are correlated with epigenetic and transcriptional changes seen in HD. These results provide, for the first time, a genome-wide map of HTT genomic occupancy and suggest that altered transcription in HD arises, in part, via alterations in direct HTT-chromatin interactions.

## Results

### HTT Reproducibly Occupies Thousands of Locations in the Mouse Genome

We set out to map the genomic occupancy of HTT and assess its relationship to HD mutations in the striatum of *Htt^Q111/Q111^* and wildtype mice. *Htt^Q111^* is a well-characterized, genetically precise knock-in mouse model of a mutation associated with juvenile-onset HD, in which a human allele of *HTT* exon 1 with approximately 111 glutamine-encoding CAG repeats has been inserted into the endogenous mouse *Htt* locus. For maximal fidelity to genetics of human HD patients, most studies utilize heterozygous *Htt^Q111/+^*mice.

However, for HTT ChIP-seq experiments homozygous *Htt^Q111/Q111^*mice are preferred to avoid the confound of two isoforms of HTT being present in each sample. HD is inherited in a fully dominant fashion, and HD patients with compound heterozygous HD mutations generally experience symptoms equivalent to heterozygous patients with a single mutant allele. Likewise, the progression of disease-related phenotypes is comparable in heterozygous *Htt^Q111/+^* mice and *Htt^Q111/Q111^*homozygotes, with four months of age representing an early time point. At this age, we[8,21] and others[10] have detected hundreds of differentially expressed genes in striatal tissue and misfolded HTT isoforms in the nuclei of many striatal SPNs from *Htt^Q111/+^*mice, but there is not yet any discernible striatal cell death or glial proliferation[21]. Chromatin immunoprecipitation and deep sequencing (ChIP-seq) was performed using striatal tissue from four-month-old *Htt^Q111/Q111^* and wildtype mice, using a well-validated antibody, EPR5526[22], which recognizes an N-terminal epitope of the HTT protein with no known differences in affinity for wildtype vs. mutant HTT isoforms. We sequenced three biological replicates from *Htt^Q111/Q111^* mice and three from *Htt^+/+^* mice, with each biological replicate consisting of pooled striata from three mice (Fig. 1A).

**Figure 1:**
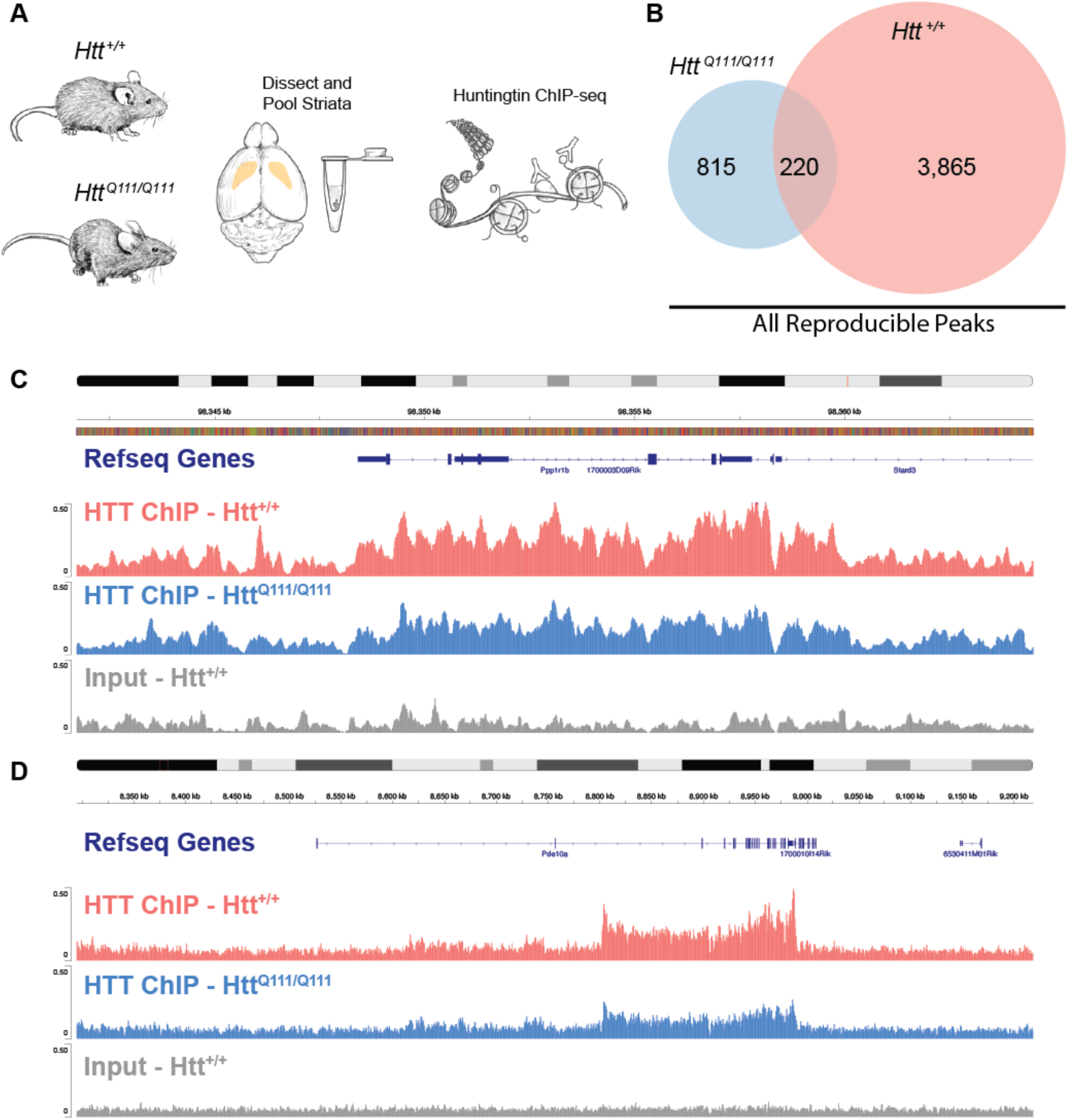
Reproducible huntingtin ChIP-seq peak profiles in 4-month-old *Htt^+/+^* and *Htt^Q111/Q111^* mice. A) Schematic of experiment. B) Venn diagram depicting the number *Htt^+/+^*-specific, *Htt^Q111/Q111^*-specific, and shared HTT peaks reproducible at FDR < 0.01 in at least 2 biological replicates from each genotype. C-D) Normalized genomic coverage at representative HTT ChIP-Seq peaks in *Htt^+/+^* mice (peach) and *Htt^Q111/Q111^* mice (blue), with *Htt^+/+^* input control sample (gray). Chromosomal locations and Refseq genes (mouse genome MM10) are indicated. Genes upstream of Pde10a in D) have been omitted for labeling clarity.

Initial inspection suggested that HTT occupies broad domains along chromatin DNA (e.g. Fig. 1C-D). We conducted peak-calling using `MACS`[23] in broad peak mode, which revealed 9,656 reproducible peaks, of which 9,624 peaks were not overlapping blacklisted regions and 4,900 peaks were conditionally reproducible, i.e. identified independently in at least two biological replicates from each genotype (Fig. 1B; Tables S1 and S2). Reproducible HTT peaks (HTT, mHTT, and shared) were found in both genic and intergenic regions: 0.1% in the 5’ UTR, 31% within 3kb upstream of TSS, 1.1% in the first exon, 3.6% in other exons, 10.7% in the first intron, 28.1% in other introns, 4.4% in the 3’ UTR, with 20.8% ≤ 300kb downstream or in distal intergenic regions (Fig. 4A). A HTT-sequence-specific control experiment supports the hypothesis that these regions accurately reflect HTT occupancy rather than non-specific signals. Namely, we observe fewer HTT ChIP-seq peaks in brain tissue from mice treated with an antisense oligonucleotide (ASO) to reduce HTT levels in the brain (Fig. S1).

Robust HTT peaks were found within 1 Mb of 2,628 protein-coding genes. Importantly, most genes do not contain HTT binding sites, suggesting that HTT occupancy in the vicinity of a given gene may reflect a specific regulatory role limited to this subset of genes, rather than promiscuous binding to every gene. Manual examination of the HTT peaks in gene bodies shows greater HTT occupancy near the 3’ end of the coding region, rather than binding as a sharp peak in a promoter or enhancer, as would be expected if HTT acts as a transcription factor[14]. Genes with this HTT occupancy pattern include canonical markers for SPNs such as *Ppp1r1b* (DARPP32) and *Pde10a* (Fig. 1C-D), which has previously been shown to have altered chromatin conformation in *Htt^Q140/Q140^* striatum[7].

### Differential Occupancy in *Htt^Q111/Q111^* mice

Next, we quantified HTT occupancy in *Htt^Q111/Q111^* vs. wildtype mice using `DiffBind`[25]. We found 244 peaks with significantly lower occupancy in *Htt^Q111/Q111^*compared to wildtype mice, and 4 peaks with higher occupancy in *Htt^Q111/Q111^* mice (FDR < 0.1; Fig. 2A; Supplementary Table S3), suggesting a bias towards reduced occupancy in *Htt^Q111/Q111^* samples. Notably, genes with nearby reduced peak occupancy in *Htt^Q111/Q111^* striata include important SPN identity genes - many with robust HTT ChIP-seq signal across the coding region - such as *Pde10a* and *Rgs9* (Fig. 2A). To more formally establish the gene sets associated with genes near `DiffBind`-nominated peaks, we assigned peaks to the nearest annotated gene if they overlapped the TSS (-1kb to +100bp), TTS (-100 bp to +1kb), exon, intronic, or intergenic regions (1kb to 1Mb), and conducted enrichment analyses using `enrichR`[26]. Genes near HTT peaks that are reduced in *Htt^Q111/Q111^* mice relative to *Htt^+/+^* mice were more likely to be markers of striatal identity, as cataloged by the Mouse Gene Atlas[27] (Fig. 2B; Supplementary Table S4). No such enrichments were found for genes near HTT peaks that are higher in the *Htt^Q111/Q111^* mice relative to *Htt^+/+^* mice (data not shown). Next, we investigated enrichments in a collection of 2,579 manually curated HD-relevant transcriptional signatures from the Huntington’s Disease Molecular Signatures Database (HDSigDB; https://hdsigdb.hdinhd.org/, 2021 mouse version). Genes near HTT peaks with decreased occupancy in *Htt^Q111/Q111^* mice were markedly enriched amongst many gene sets from HD mouse studies (Fig. 2C). Most notably, these included genes that are downregulated in the striata of preclinical HD mouse models. These data suggest a pattern of reduced HTT genomic occupancy near SPN identity genes that are transcriptionally downregulated in the striatum across HD model systems.

**Figure 2:**
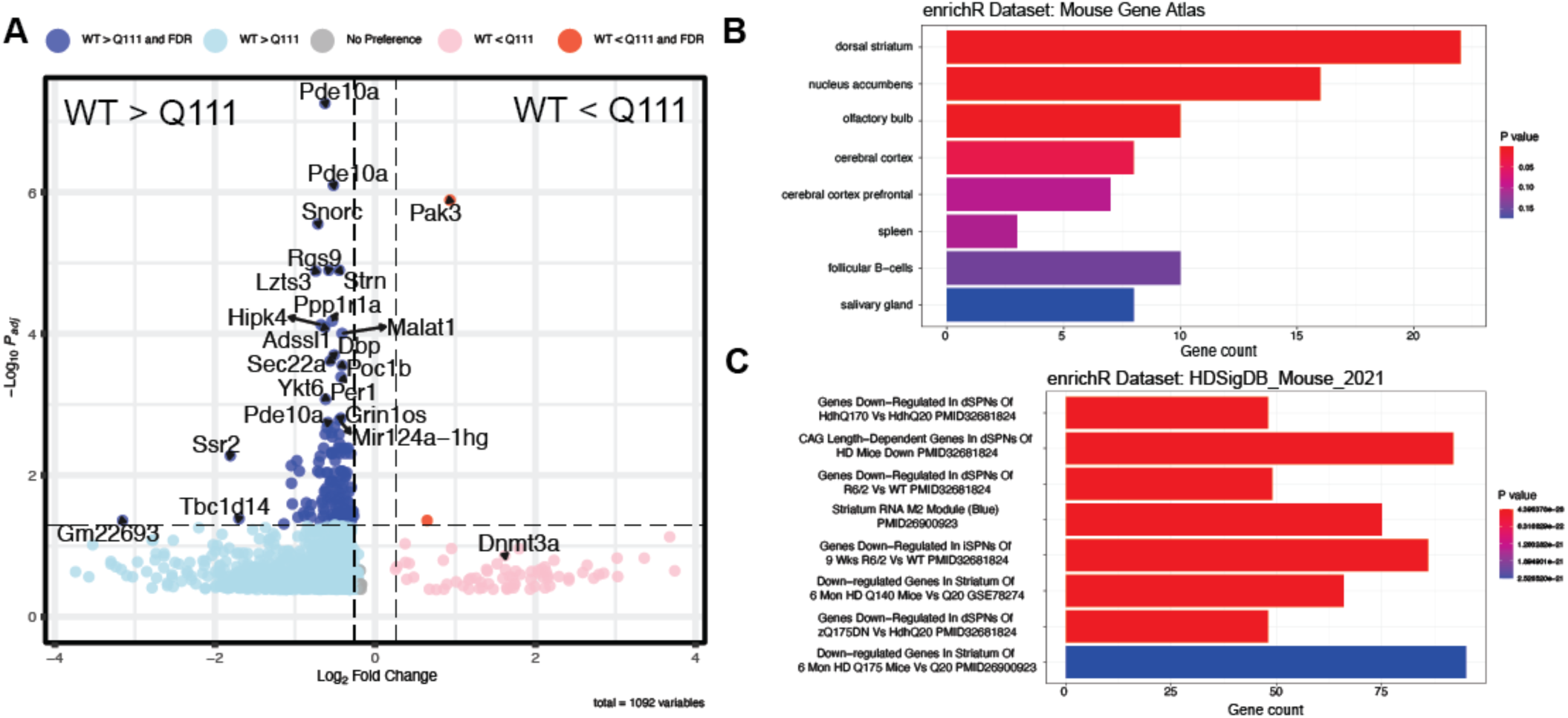
Differential chromatin binding by Huntingtin in *Htt^+/+^* and *Htt^Q111/Q111^*mice. A) Volcano plot summarizes the results of the `DiffBind` analysis – the X-axis indicates the fold change and the y-axis the adjusted *p*-value. `DiffBind` peaks for which the FDR is less than 0.05 and/or the absolute fold-change is > 0.26 are identified by the nearest gene symbol. `DiffBind` peaks not matching those criteria are indicated with gray dots. B) 244 genes with reduced peaks in the *Htt^Q111/Q111^* mice (FDR < 0.1) were examined using `enrichR` for enrichment amongst a number of different mouse tissues. The bar graph indicates the results of these genes amongst mouse tissues with significant enrichments. Only the terms “dorsal striatum” and “nucleus accumbens” meet the stringency threshold *P_adj_* < 0.05. C) Enrichment of `DiffBind` nominated peaks in HDSigDB gene sets reveals strikingly enriched sets, notably those that include genes which are down-regulated in the striatum of HD mouse models.

### Histone Methylation and EZH2 Occupancy in *Htt^Q111/+^* Mice

To compare HTT occupancy and chromatin states of interest, we generated ChIP-seq of Histone H3 modifications associated with specific chromatin states: trimethylation of lysine 27 (H3K27me3), trimethylation of lysine 9 (H3K9me3), trimethylation of lysine 4 (H3K4me3) and acetylation of lysine 27 (H3K27ac). These marks are associated with active promoters (H3K4me3), active promoters and enhancers (H3K27ac), facultatively repressed promoters and enhancers (H3K27me3), and constitutively repressed regions (H3K9me3). These experiments used striatal tissue from mice of the same age as the *Htt^Q111/Q111^* animals used in our HTT ChIP-seq experiments described above (N=3/genotype), but utilized heterozygous *Htt^Q111/+^*mice to enable matched comparisons to published gene expression datasets from heterozygous *Htt^Q111/+^* mice[8,10,21]. After library construction, sequencing, input normalization and peak calling, we identified 17672, 25315, 22304, and 27337 reproducible peaks for H3K4me3, H3K27me3, H3K9me3, and H3K27ac, respectively. As expected, H3K4me3 was localized primarily at promoters, H3K27ac and H3K27me3 in enhancers and promoters, and H3K9me3 in more distal regions (Fig. 4A). Using these data, we examined differential methylation, acetylation or occupancy of each dataset in *Htt^Q111/+^* vs. *Htt^+/+^* samples. At FDR < 0.1, there were no robust peak differences for H3K9me3, H3K4me3, or H3K27ac, suggesting that any genotypic differences in these marks are subtle. However, we identified 42 H3K27me3 differentially methylated regions (DMRs) at FDR < 0.1 (Supplementary Table S5; Fig. 3A). Notably, 41 of 42 H3K27me3 DMRs (98%) had reduced levels of methylation in *Htt^Q111/+^* vs. *Htt^+/+^*. At a more lenient p-value (P < 0.05), there were 2018 H3K27me3 DMRs, of which 1926 (96%) had reduced methylation in *Htt^Q111/+^* samples. These results suggest that HD mutations lead to changes in H3K27 trimethylation in the 4-month-old *Htt^Q111/+^*striatum, including a previously undescribed reduction in H3K27me3 levels in many genomic regions.

**Figure 3.**
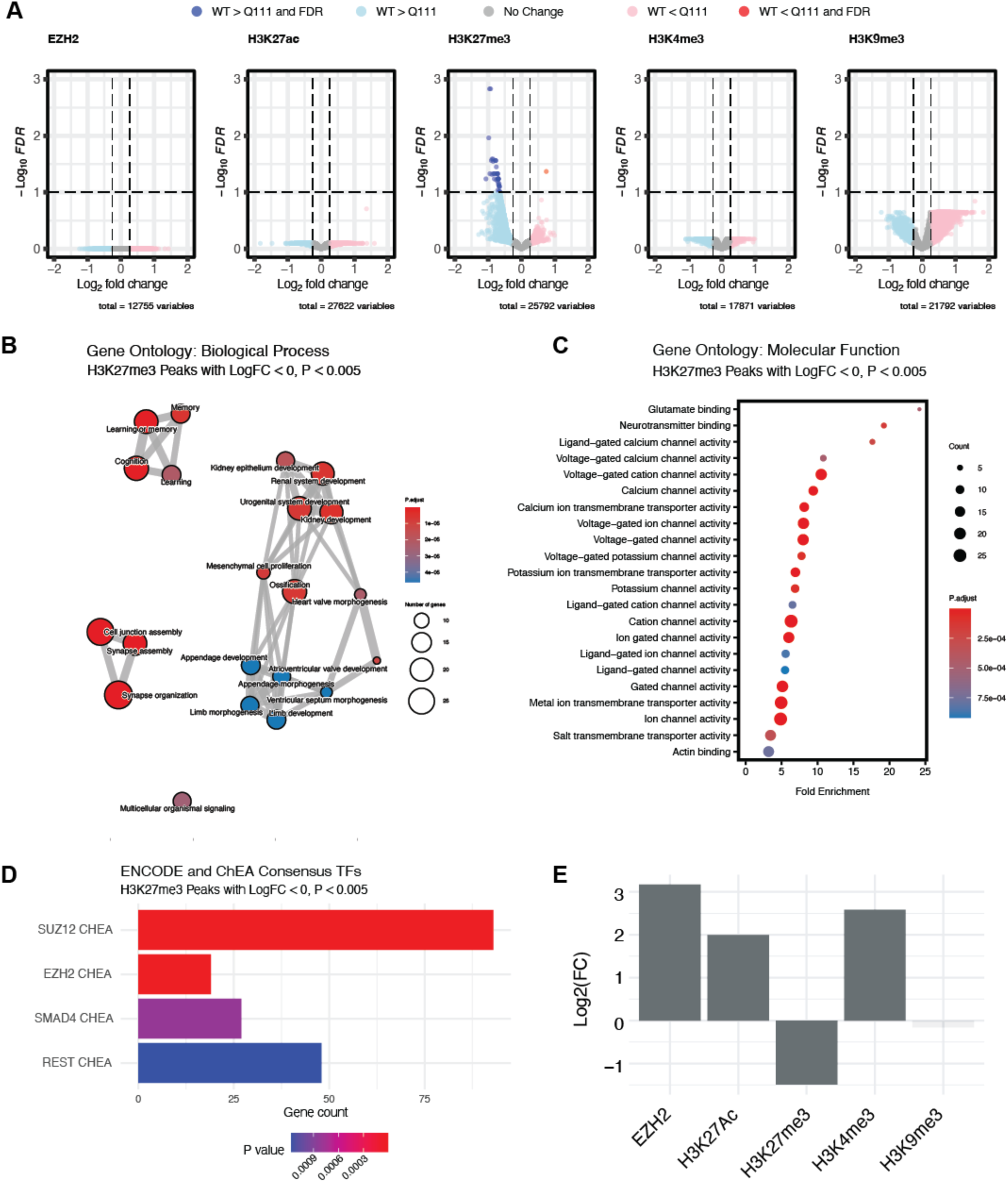
ChIP-seq analysis reveals early changes in H3K27me3 near developmentally important genes in *Htt^Q111/+^* mice. A: Volcano plots summarizing differential histone modifications for H3K27me3, H3K9me3 and H3K4me3 and differential occupancy for EZH2 in striatal tissue from four-month-old in *Htt^Q111/+^* vs. *Htt^+/+^* mice. The x-axis indicates the log_2_FC and the y-axis indicates the -log_10_(p-value) for the comparison of peak read depth between genotypes. Regions with greater levels or occupancy in the *Htt^+/+^*mice are shown in blue, regions with greater levels or occupancy in the *Htt^Q111/+^*mice are shown in pink/red. The dashed horizontal line indicates an FDR of 0.1 and vertical lines are Log_2_FC = +/-0.26. B) Network depiction of Gene Ontology Biological Processes enrichment near differentially methylated regions with reduced H3K27me3 in *Htt^Q111/+^* mice. C) Gene Ontology Molecular Function enrichment near differentially methylated regions with reduced H3K27me3 in *Htt^Q111/+^* mice. D) Enrichment of 495 genes in nominally differentially methylated regions with reduced H3K27me3 in *Htt^Q111/+^* mice (unadjusted p < 0.005) amongst a consensus transcription factor/target gene databases (i.e. ENCODE and ChEA Consensus TFs from the `enrichR` package). Robustly enriched transcription factor lists include: SUZ12 CHEA (93/1,684; *p_adj_* = 8.3e13; EZH2 CHEA (19/237; *p_adj_* = 2.0e04); SMAD4 CHEA (27/584; *p_adj_* = 2.6E02); REST CHEA (48/1,280; *p_adj_* = 2.8e02). E) Enrichment of EZH2 and indicated chromatin mark ChIP-seq peaks (MACS FDR < 0.05) in huntingtin ChIP-seq peak regions. Y-axis indicates the log-transformed fold change (enrichment or depletion) in the number of overlapping base pairs compared to the average from 100,000 re-sampling permutations of genomic coordinates. Plotting color indicates the p-value, derived from these same permutations.

**Figure 4.**
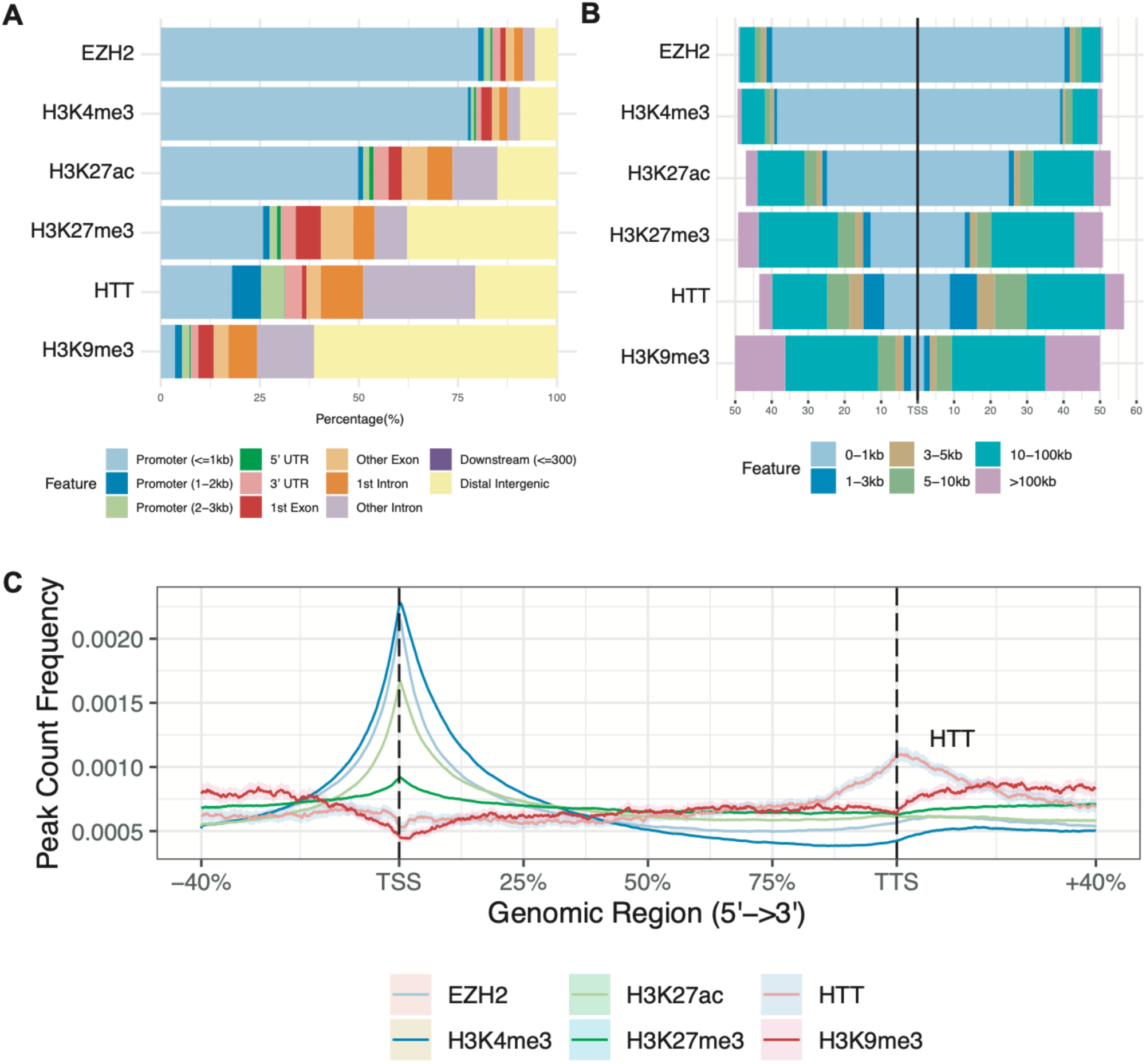
Huntingtin peaks occupy specific chromatin functional regions. A) Summary of genomic regions occupied by huntingtin and other histone mark / EHZ2 occupancy peak sets generated in this study. The percentage of peaks for each dataset are indicated on each vertical line, with boxes indicating the percentage of that marks found in each of the categories indicated. B) Localization of peaks for the indicated chromatin-associated protein at binned distanced from an averaged transcriptional start site (TSS). C) Peak count frequencies of genic ChIP-seq peaks across the normalized transcript length of all mouse transcripts. The average peak count is indicated via the black bar, from 5’ (left) to 3’ (right). Genomic regions are indicated as percentages, with 100% spanning from the TSS to the transcriptional termination site (TTS). Shading indicates bootstrapped 95% confidence intervals calculated with resampling (ChIPSeeker package).

The reduction of H3K27me3 in *Htt^Q111/+^* vs. *Htt^+/+^*striata suggested a possible partial change of PRC2 function, the enzymatic complex responsible for “writing” the H3K27me3 mark[20]. We investigated the genes proximal to H3K27me3 DMRs (p-value < 0.05) to gain insight into their potential biological consequences.

Functional annotation of H3K27me3 DMRs with `GREAT` [28] and `Genekitr` [29](Fig. 3B; Supplementary Table S6) revealed that negative DMRs in *Htt^Q111/+^* striata were enriched near genes that impinge on development and morphogenesis (e.g. GO:0001822, Kidney Development, *p_adj_* = 4.4e-06), genes involved in synapse formation and stability (e.g. GO:0007416, Synapse Assembly, *p_adj_* = 1.4e-10), and genes related to cognitive processes (e.g. GO:0007611, Learning or Memory, *p_adj_* = 5.3e-07). Genes associated with decreased H3K27me3 levels were enriched for molecular functions at the synapse (Fig. 3C), including glutamate binding (GO:0016595, *p_adj_* = 4.8e-04) and calcium channel activity (GO:0005262, *p_adj_* = 9.7e-08). Assessment of negative H3K27me3 DMRs for transcription factor-associated chromosomal localization using `enrichR` indicated enrichment at known PRC2 target genes (i.e. those associated with PRC2 subunits EZH2 and SUZ12; Fig. 3D), many of which are involved in cell fate determination[30]. Notable examples of developmentally important genes near negative H3K27me3 DMRs include the TFs *Sox11* and *Foxp2.* Therefore, mHTT expression in the striatum is associated with reduced H3K27me3 levels in the vicinity of synaptic, as well as developmentally important genes, including transcription factors that are targets of the PRC2 complex. To further explore this, we performed ChIP-seq of EZH2, the enzymatic component of PRC2 that is responsible for methylation of H3K27, in the striatum of four-month-old *Htt^Q111/+^* and wildtype mice (n = 3 / genotype), identifying 12388 reproducible peaks. Interestingly, this analysis revealed no differentially occupied sites (Fig. 3A), suggesting that HD does not alter the localization of the EZH2-containing PRC2 complex itself at this age.

We next tested the hypothesis that HTT ChIP-seq peaks co-localize with histone modifications and EZH2 (Fig. 3E; Supplementary Table S7). We found a striking overlap between our HTT peaks and EZH2 (log_2_FC = 3.2; *p_adj_* = 3.2e-05), H3K27ac (log_2_FC = 2.0; *p_adj_* = 3.2e-05), and H3K4me3 (log_2_FC = 2.6; *p_adj_* = 3.2e-05). We did not observe global overlap between constitutive heterochromatin marked by histone H3 lysine 9 trimethylation (H3K9me3; log_2_FC = -0.2; *p_adj_* = 0.41), and we observed a *depletion* of HTT peaks in H3K27me3 peak regions (log_2_FC = -1.5; *p_adj_* = 3.2e-05). This apparent discrepancy between HTT peaks being enriched in EZH2 regions, but depleted from H3K27me3 peak regions, may indicate differences in the association of HTT with sites of active EZH2-containing PRC2 complexes versus stably methylated H3K27.

### HTT Binds Specific Chromatin Domains

To explore a role for HTT in gene regulation, we compared the locations of all 9624 HTT peaks we observed to known genomic features using the `ChIPseeker` Bioconductor package[24]. HTT’s occupied genomic regions differed substantially compared to the other marks (Fig. 4A), with a notable enrichment within introns, consistent with HTT’s consistent coverage over coding regions of target genes. Considering the abundance of all peaks relative to the transcriptional start site (TSS), huntingtin peaks were substantially less enriched near the TSS compared to marks canonically associated with active transcription and open chromatin (H3K4me3, H3K27ac) and those of EZH2, which also occupied regions upstream of the TSS of target genes, and less enriched in distal intergenic regions than H3K9me3 (Fig. 4B). HTT peaks had a distinct profile relative to the TSS compared to each of the other mark examined (Fig. 4B), and of the marks surveyed here were the only ones with increased signal near the transcription termination site (TTS; Fig. 4C).

### HTT occupancy predicts effects gene expression in HD

The selective enrichment of HTT binding within regions of chromatin bearing marks associated with active transcription (H3K4me3 and H3K27ac; Fig. 1E), and its depletion from a mark associated with repressive transcription (H3K27me3; *ibid*) raised the possibility that HTT may directly contribute to HD-related changes in gene expression. We mapped potential HTT target genes whose TSSs were located within +/- 20 kb of a HTT ChIP peak. Assigning regulatory elements to their target genes is inevitably inexact, but previous work has shown that summing the regulatory elements within 20 kb of a TSS optimizes sensitivity and specificity for detecting gene regulatory interactions[31]. Given HTT’s striking occupancy at specific genes (e.g. Fig. 1C-D, Fig. 2), we considered whether it might more generally occupy regions near genes with disease-relevant changes in gene expression. We first considered the simplest category of interest: genes that are up- or down-regulated in the striatum of HD model mice, with a focus on the *Htt^Q111/+^* striatum at 6 months of age, taken from a larger allelic series study[10]. We find enrichment of HTT peaks (irrespective of peak category, or “All Reproducible”) near genes that are downregulated in the striatum of 6-month-old *Htt^Q111/+^* mice (Fig. 5A; log_2_FC = 0.73, *p_adj_* = 1.4e-30) and depleted near genes that are up-regulated (Fig. 5A; log_2_FC = -0.33, *p_adj_* = 5.3e-05). Expanding this analysis to differentially expressed genes in other mouse lines and timepoints from the allelic series study reinforced our observation that HTT-occupied chromatin peaks are more likely to be associated with down-regulated genes than up-regulated genes (Fig. 5B, Supplementary Table S8).

**Figure 5.**
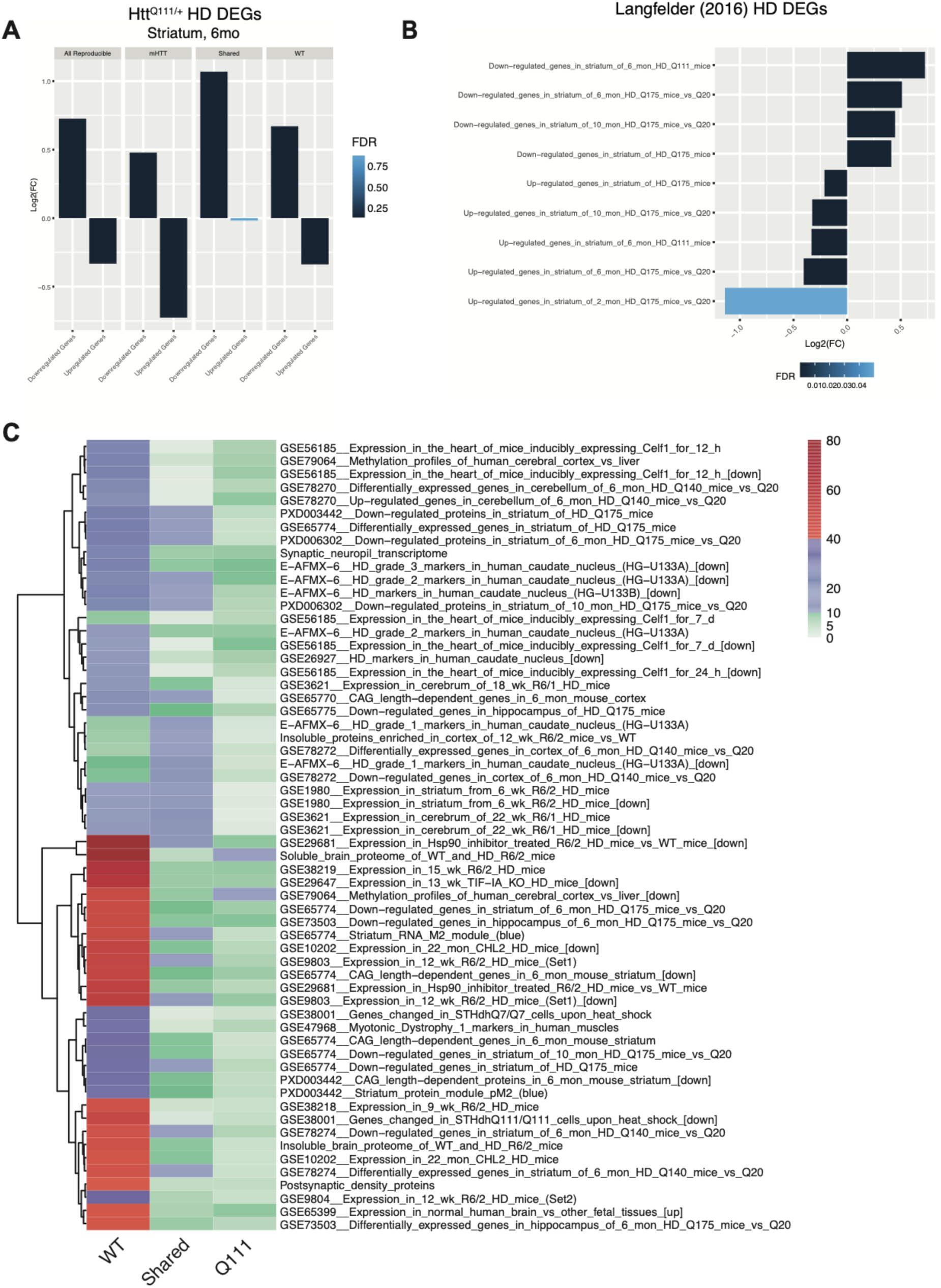
Genotype-specific HTT occupancy in genomic regions surrounding HD DEGs and other CAG-sensitive gene sets of interest. A-B) Enrichment or depletion of HTT ChIP-seq peaks in the regions +/- 20 kb from the transcription start sites of up- and down-regulated differentially expressed genes (DEGs) in the striatum of knock-in mouse models of HD mutations, based on RNA-seq from [10,46]. Y-axis indicates fold enrichment of base pair overlap compared to the average from 100,000 re-sampling permutations. C) Heatmap showing HDSigDB gene sets enriched in the regions +/- 20 kb of WT-specific, Q111-specific, and Shared HTT ChIP-seq peaks. The plotting color indicates the -log_10_(p-value) for the strength of the enrichment, based on 100,000 re-sampling permutations. Rows are ordered by hierarchical clustering, based on p-values.

We next examined the enrichment of our HTT peaks amongst all manually curated gene sets cataloged in the HDSigDB database (Fig. 5C, Supplementary Table S9). As with our `DiffBind`-nominated peaks with differential binding in *Htt^Q111/Q111^* versus *Htt^+/+^*mice (Fig. 2), this analysis revealed a striking enrichment amongst genes that are dysregulated in a number of studies, and particularly in striatal genes that are down-regulated in HD (e.g. Fig. 5B). A particularly notable gene set is the “Striatum RNA M2 Module (blue)” from an exhaustive molecular network characterization of an allelic series of HD knock-in mice[10]. This module of genes is notable for being tightly correlated with CAG length and age in the striatum of mHTT-expressing mice and contains many striatal identity genes. This suggests that our HTT peaks are markedly enriched amongst striatum-expressed genes, including SPN identity genes which are down-regulated in HD. Given PRC2’s role in regulating cell-specific gene expression and described interactions with HTT[17], we specifically focused on PRC2-relevant gene sets in HDSigDB. Our shared HTT peaks, those found in both wildtype and Q111/Q111 mice, were strikingly enriched in bivalent genes whose expression is increased at 3- (Fig. 5C; Odds ratio = 16.7, FDR = 7.6e-05) and 6-months of age (Fig. 5C; Odds ratio = 6.2, FDR = 3.6e-03) in the striatum of mice lacking PRC2 expression in SPNs (*Ezh1^-/-^;Ezh2^-/-^*)[18]. This suggests that our peaks are near genes that are both down-regulated in HD, *and* up-regulated in the context of PRC2 knockout, supporting the hypothesis that HTT binding at a small set of bivalent genes may play an important role in gene dysregulation in HD. Consistent with our own data (Fig. 3D), revealing a depletion of HTT peaks in H3K27me3 regions, our mutant-specific HTT peaks were depleted near H3K27me3-enriched genes in purified SPNs (Fig. 5C; Odds ratio = 16.7, FDR = 7.6e-05)[18] .

### Correlations Between HTT Binding and Epigenetic Features Near HD DEGs

We hypothesized that the negative correlation between HTT occupancy and HD-related gene expression may be mediated by PRC2 and histone modifications. To investigate this, we integrated our data on HTT occupancy, histone modifications, and transcription, focusing on the genomic regions proximal to DEGs in 6-month-old *Htt^Q111/+^* versus *Htt^Q20/+^* mice (Fig. 6A-C; Table S8 [10]). We defined intervals of interest +/- 20 kb of the TSS of each HD DEG and calculated the fold change in *Htt^Q111/Q111^*vs. *Htt^+/+^* mice for HTT, H3K27me3, H3K4me3, and EZH2 occupancy within the same interval. Consistent with our hypothesis, the fold changes in H3K27me3 were constitutively lower and negatively correlated with HTT occupancy near genes differentially expressed in HD (Fig 6B; Spearman’s rho = -0.22; p = 1.3e-3). This suggests that *Htt^Q111/+^* mice have generally lower H3K27me3 levels near HD DEGs, and that increased HTT binding in a region is associated with reduced H3K27me3 in the same interval. Conversely, in those same regions we observe constitutively higher, and positively correlated, relationships across genotypes between HTT binding and H3K4me3 (Fig 6C; Spearman’s rho = 0.27; p < 1e-300) and EZH2 (Fig. 6A; Spearman’s rho = 0.14; p = 1.5e-2). That is, in *Htt^Q111/+^* mice, near HD DEGs, levels of EZH2 occupancy and H3K4me3 were generally higher, with increased HTT binding in that region associated with more EHZ2 occupancy and higher H3K4me3 levels in that same interval. Conversely, levels of H3K27me3 near HD DEGs were lower in Htt^Q*111/+*^ mice, and greater local HTT binding predicts lower H3K27me3. Importantly, these relationships appeared to hold true both for up- and down-regulated genes in *Htt^Q111/+^* mice. Thus, concordant changes in PRC2 localization and histone methylation/demethylation may coincide with alterations in HTT occupancy in these regions in both *Htt^Q111/+^* and *Htt^+/+^* mice.

**Figure 6.**
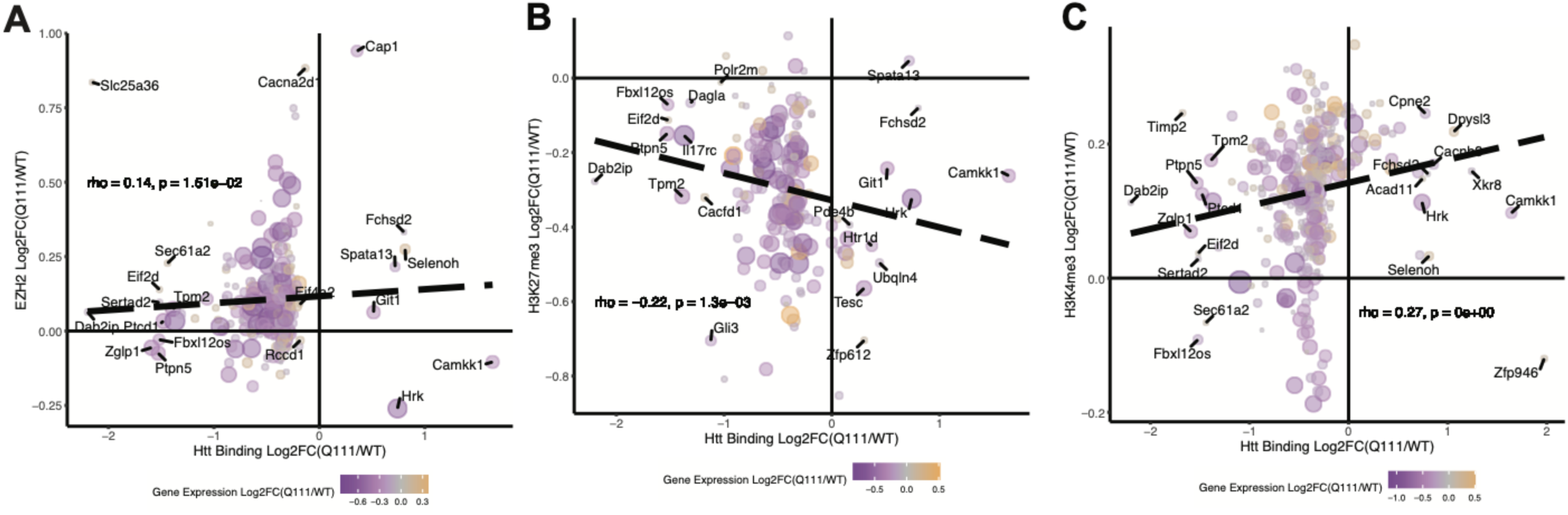
Increased HTT occupancy is negatively associated with H3K27me3 and positively associated with H3K4me3, near genes that are dysregulated in HD. Each plot indicates the fold change of HTT occupancy in *Htt^Q111/Q111^* vs. *Htt^+/+^* mice (x-axis) compared to the fold change in the same regions for EZH2 (A), H3K27me3 (B), and H3K4me3 (C) on the y-axis. Analysis is restricted to 263 HTT peaks located +/- 20 kb from the TSSs of genes that are significantly up- or down-regulated in the striatum of six-month-old knock-in mouse models of HD mutations. Each point represents a single HTT ChIP-seq peak. Point color indicates the fold change in gene expression in the striatum of HD knock-in mice (up = yellow; down = purple), and point size corresponds to the p-value for differential expression.

## Discussion

This study was motivated by the question of whether transcriptional dysregulation in Huntington’s disease could occur due to interactions between HTT and chromatin. Previously, we investigated whether specific transcription factors demonstrated differential genomic occupancy and compared this with gene expression changes in *Htt^Q111/+^* mice, finding that the transcription factor SMAD3 demonstrated differential binding between wildtype and mutant mouse brain tissue[10,32]. This finding led us to ask what role the HTT protein itself might play with regards to codified chromatin interactions.

ChIP-seq of the HTT protein reveals thousands of robust and reproducible sites of HTT genomic occupancy in the mouse striatum, the brain region most vulnerable to HD pathology and in which the most pronounced transcriptional dysregulation occurs. Many HTT peaks are enriched across the coding sequence of genes of particular interest in HD, with a surprising enrichment near the 3’ end of many transcripts. We further observed, for the first time, a reduction in the levels of H3K27me3 in the striatum of young *Htt^Q111/+^*mice, suggesting these changes represent a very early stage of chromatin alterations in the most vulnerable brain region in HD, which experiences the greatest burden of transcriptional changes in HD. HTT peaks are enriched near genes that are down-regulated in HD, and depleted near genes that are up-regulated, supporting a link between these HTT:chromatin interactions and bi-directional gene expression changes in HD. An integrated analysis of HTT binding and histone marks suggests that local HTT binding near HD DEGs is associated with increased H3K4me3 and EZH2 binding, and reduced H3K27me3 levels.

HTT physically interacts with transcriptional regulatory proteins such as p53 and CREB-binding protein[33]. In addition, previous ChIP-qPCR experiments with HTT antibodies suggested occupancy in promoter regions of specific genes[14]. With the improved resolution of genome-wide ChIP-seq, we observed thousands of robust HTT:chromatin interactions in a specific subset of genes. However, these HTT peaks are not primarily localized to the promoters. Instead, distributions of HTT occupancy across genic regions were more reminiscent of marks such as H3K36me3[34,35] and RNAPII Ser2[36,37] that are associated with transcriptional elongation and enriched at the 3’ end of the gene, a pattern distinct from active marks such as H3K4me3 that are more strongly associated with TSSs[38,39]. Thus, both the gene identity and pattern of HTT binding in those genes argues for a specific, selective, role for HTT-chromatin interactions. This argument is strengthened by the fact that the specific genes with robust HTT occupancy in the striatal tissue we profiled are enriched for SPN identity genes which we [19], and others[10,19], have found to be strikingly downregulated in HD.

Many genes with HTT peaks identified in our study are targets of the PRC2 complex, a critical regulator of cell identity via regulation of the H3K27me3 mark near genes important for cell-fate commitment[20], including in SPNs of the striatum[18], the most vulnerable brain region to HD pathology[2,18]. Direct interactions between HTT and the PRC2 complex have been described, and HTT has been proposed to directly enhance the activity of the enzymatic activity of the PRC2 complex[17]. The enrichment of HTT within well-annotated PRC2 target genes supports a role for HTT in regulating their expression, however, in the aggregate, the occupancy patterns of EZH2 and HTT are quite distinct (Fig. 4A-C). And while, genome-wide, we observe robust overlap between HTT and EZH2 peaks, we observe a *depletion* of HTT in H3K27me3 peak regions (Fig. 3D), suggesting HTT is unlikely to occupy all chromatin-bound PRC2 complexes, nor remain associated with facultative heterochromatin, once formed. Given its role in scaffolding functional protein-protein interactions[17,40], and dynamic shuttling between the nucleus and cytoplasm[41], we hypothesize that HTT may play a role in regulating the balance between active and repressed chromatin at select PRC2 target genes.

Several potential mechanisms for HTT’s involvement in transcriptional regulation via the PRC2 complex deserve attention in future studies. A functionally important role for interactions between HTT and the repressor element 1 silencing transcription factor (REST), which plays important roles in repressing neuronal gene expression in non-neuronal cells, have been described[42]. Namely, HTT and REST bind, and HTT aids in confining REST to the cytoplasm, thereby preventing aberrant expression of REST target genes in non-neuronal cells. Expression of mHTT results in the aberrant accumulation of REST in the nucleus, and consequential dysregulation of REST-mediated signaling. With this background, we were intrigued with our results demonstrating that regions with lower H3K27me3 in the striata of *Htt^Q111/+^*mice are enriched in both PRC2 and REST target genes (Fig. 3D), especially considering data suggesting that REST may be involved in targeting PRC2 to specific loci[42,43]. Recently, a new pathway for regulation of PRC2 function via cytoplasmic retention of embryonic ectoderm development (EED), a PRC2 subunit critical for formation of the complete trimeric structure, has been described[44]. In post-mitotic myotubes, a short cytoplasmic form of PRC2 subunit EZH1 (EZH1β) sequesters EED in the cytoplasm, preventing formation of the PRC2 complex at target genes in the nucleus. Future work focused on understanding whether HTT plays a role in the regulation of PRC2 signaling via cytoplasmic sequestration of PRC2 subunits may be a fruitful area of investigation.

We complemented our HTT ChIP-seq findings with ChIP-seq of histone modifications and EZH2 binding, which revealed reductions in H3K27me3 in the striatum of young *Htt^Q111/+^* mice, similar to those seen in *Htt^Q140/+^* mice[7] and particularly near important striatal cell identity genes (e.g. *Pde10a*). Our differential acetylation data do not recapitulate large-scale genotype-dependent changes in H3K27ac previously seen in HD mouse model striatum, which could be due to our use of bulk tissue versus fluorescence-activated nuclear sorted neurons[7]. Further, while we do see association of HTT peaks with H3K27ac (Fig. 4E), HTT shows less association than with EZH2 and H3K4me3, suggesting that HTT is likely not localized to super-enhancers but instead to gene bodies (Fig. 3A). The observed reduction in H3K27me3 in *Htt^Q111/+^* mice is consistent with a hypothesis of PRC2 complex dysfunction and/or trafficking disruption occurring at the earliest stages of progression in this model, though other chromatin modifiers could also play roles. The identified lower levels of H3K27me3 seen in heterozygous polyQ HTT-expressing striatum and lack of changes in EZH2 occupancy in particular may implicate H3K27-targeted histone demethylases such as KDM6A/UTX in epigenetic changes observed in HD[45]. While correlational, our integrated analysis of HTT, EZH2 binding, and histone marks suggests that increased HTT binding near HD DEGs in *Htt^Q111/+^* mice predicts increased H3K4me3 and EZH2 binding, with reduced H3K27me3 at these same loci. We believe detailed mechanistic experiments focused on elucidating the pathways linking these events could provide important new insights into HTT’s role in transcriptional dysregulation in HD.

This study has several important limitations. Like all ChIP-seq studies, peak detection is sensitive to background noise, antibody cross-reactivity, and other sources of bias. We have tried to control for this by using a relatively large number of technical and biological replicates. Another consideration is the reliance on antibodies, which limits us to establishing chromatin regions associated with specific epitopes of HTT. Future work using phospho-specific antibodies, and other dynamic epitopes is likely to nominate additional HTT-chromatin interactions, which are of great interest. Functional enrichment analyses of our diffbind results from wildype HTT versus mHTT rely on gene assignment that may be biased towards longer genes, propagating these biases through the enrichment tools. This is likely less of an issue for HTT, since many of its peaks are directly over gene bodies, but highlights that predicting the potential regulatory functions of distal peaks is difficult. Finally, our ChIP-seq experiments relied on bulk striatal tissue, whereas single-nuclei RNA sequencing has indicated bidirectional effects of HD mutations across cell types, especially for PRC2 targets[19]. Emerging techniques enabling ChIP-seq of single cells and sorted cell populations[19,46] should enable additional refinements of the findings presented here.

While ChIP-on-chip has been performed to identify HTT binding sites at promoter sequences[14], this work provides the first genome-wide map of HTT-chromatin interactions and has identified key changes in these interactions due to HD mutations. It suggests that aberrant de-repression of CAG-sensitive genes in HD samples–including cell identity genes–may be due to molecular interactions of HTT and chromatin. We find that altered HTT-chromatin occupancy is accompanied by novel histone modification changes – notably, reductions in H3K27me3 – in the *Htt^Q111/+^* striata, which are associated with genotype-selective HTT occupancy in the same regions. In fact, the strength of HTT occupancy predicts increased H3K4me3 and EZH2 occupancy, and reductions in H3K27me3, consistent with a model in which HTT plays a functional role in regulating PRC2 activity, perhaps by helping recruit PRC2 to specific loci.

### Conclusions

Collectively, our data support a model of pathogenesis in which perturbation of normal HTT chromatin regulatory functions precipitates transcriptional dysregulation in HD, leading to epigenetic changes and loss of cell type-specific gene expression.

## Materials and Methods

### Mouse Breeding, Genotyping, and microdissection

The B6.*Htt^Q111^* mice (Strain 003456; JAX) used for the ChIP-seq study have a targeted mutation replacing mouse *Htt* exon 1 with the corresponding portion of human *HTT* exon 1, including an expanded CAG tract. The targeted *Htt* allele was placed from the CD-1 background onto the C57BL/6J genetic background by selective backcrossing for more than 10 generations to the C57BL/6J strain at Jackson laboratories. Cohorts of homozygote, heterozygote and wildtype littermate mice were generated by crossing B6.*Htt^Q111/+^* and B6.*Htt^+/+^* mice. Male mice were sacrificed at four months of age via a sodium pentobarbital based euthanasia solution (Fatal Plus, Henry Schein). Brain tissues were snap frozen in liquid nitrogen and stored in -80°C until ChIP was performed. Experiments were performed following NIH animal care guidelines and approved by Western Washington University’s Institutional Animal Care and Use Committee under protocols 14-005 and 16-011.

### Huntingtin knockdown in cortex

Four-month-old male mice underwent unilateral intracerebroventricular injection of 500ug pan-huntingtin antisense oligonucleotide (Ionis #444652) or received no treatment. Tissue was collected as above at four weeks post-injection; a time shown to have maximal HTT reduction in BACHD mice[47]. Cortical tissue contralateral to the injection was used to assess HTT knockdown by western blot as described[48], and ipsilateral tissue was used for ChIP as performed below.

### HTT ChIP-seq

We prepared replicate (*n* = 3) ChIP samples using an anti-huntingtin antibody from four-month-old male *Htt^Q111/Q111^* and age-matched *Htt^+/+^* mice. For each ChIP preparation, chromatin DNA was prepared using the combined striatal tissue from both hemispheres of three mice. Preliminary experiments suggested that this was the minimal amount of material required to provide enough material for multiple IPs. Striata were transferred to a glass Dounce on ice and homogenized in cold phosphate-buffered saline with protease inhibitors. For ASO-treated cortices, two cortices were pooled for each replicate (n = 1). High-resolution X-ChIP-seq was performed as previously described [49], with slight modifications[32]. IPs were performed using Abcam anti-huntingtin EPR5526 (#ab109115). ChIP-seq library preparation and sequencing reactions were conducted at GENEWIZ, Inc. (South Plainfield, NJ, USA). Sequencing was performed on an Illumina HiSeq 4000 using 2x150 Paired End (PE) configuration. Sequencing reads were aligned to the mouse genome (mm10) using HISAT2. Peak-calling was then performed with MACS2.2, scaling to the size of the input control library. The final set of reproducible peaks was obtained using the `dba` function in the `DiffBind` R package[25], retaining peaks that were reproducible at FDR < 0.01 in at least two samples and excluding artifactual blacklist regions from ENCODE. Sequence reads have been deposited in gene expression omnibus, accession GSE150750.

### Histone mark and EZH2 ChIP-seq

Fresh frozen striatal tissue from five male four-month-old *Htt^Q111/+^*and age-matched *Htt^+/+^*mice were pooled per replicate for *n* = 3 samples. Further processing was performed at Active Motif (Carlsbad, CA). Tissue was fixed in 1% formaldehyde, lysed, and disrupted with a Dounce homogenizer. DNA was sonicated to an average fragment length of 300-500bp, and 25ug chromatin plus 200ng *Drosophila* spike-in chromatin was incubated with antibody targeting EZH2, H3K27me3 or H3K4me3 (Active Motif catalog numbers 39901, 39155 and 399159, respectively). Antibody against H2Av was also present in the reaction to ensure efficient pull-down of the spike-in chromatin. Complexes were captured using protein A agarose beads (Invitrogen). Illumina sequencing libraries were prepared from ChIP and Input DNA by end-polishing, dA-addition, and adaptor ligation. Libraries were quantified and sequenced on Illumina’s NextSeq 500 (75nt single end reads).

### Analysis of striatal HTT ChIP-seq data

For the primary striatal HTT ChIP-seq dataset, sequencing reads were aligned to the mouse genome (mm10) using `bowtie2` [50]. Peak-calling on each sample was performed with MACS v2.1 [23], scaling each library to the size of the input DNA sequence library to improve comparability between samples. We retained peak regions with a MACS p-value < 0.001. Filtered peak calls were concatenated across all *Htt^+/+^* and *Htt^Q111/Q111^*samples to produce a combined set of peak calls and removed peaks overlapping artifactual blacklisted regions of the genome[51].

### Analysis of cortex HTT ChIP-seq data

Sequencing reads were aligned to the mouse genome (mm10) using HISAT2. Aligned reads from the four ChIP libraries were down-sampled to the size of the smallest sequencing library using `samtools` view -s[52]. Aligned reads from the input genomic DNA of all four samples were merged into a single control BAM file. Peak-calling was then performed with MACS2.2, scaling to the size of the input control library. The final set of reproducible peaks was obtained using the `dba` function in the `DiffBind` R package, retaining peaks that were reproducible at FDR < 0.01 in at least two samples and excluding artifactual blacklist regions from ENCODE [38]. We then used `dba.count` to count the reads in each peak region from each ChIP sample and from the control sample. Next, a generalized linear model was fit, using the `dba.analyze` DESeq2 wrapper, and we performed log-ratio tests to estimate effects of ASO treatment on HTT occupancy in each peak region. Genotype was treated as a blocking factor. Control read counts were subtracted for each site in each sample before fitting the model. This experiment was underpowered to detect statistically significant changes in occupancy at individual peak regions. Therefore, our primary test was for global depletion of HTT occupancy in ASO-treated mice across peak regions. For this purpose, we computed one-sided binomial tests with `binom.test`, comparing the number of peaks with increased vs. decreased occupancy.

### Analysis of histone mark and EZH2 ChIP-seq

Reads were aligned consecutively to the mouse (mm10) and *Drosophila* (dm3) genomes using the BWA algorithm (default settings). The number of mouse alignments used in the analysis was scaled to the number of *Drosophila* spike-in alignments [53]. Peak-calling was performed with MACS v2.1. We selected peaks that were reproducibly identified in at least two samples of the same genotype. For analyses of differential occupancy, reads mapped to each peak region were normalized to total library size. Generalized linear models and log-ratio tests were fit with `edgeR`[54] to identify differentially methylated regions.

### Enrichment of peaks in genomic regions and gene sets

Over-representation of HTT peaks in chromatin states, genomic regions marked by histone modifications, and gene sets were calculated using the Genomic Association Tester (GAT; [55]). GAT calculates the number of base pairs overlapping between two genomic annotations and estimates its fold enrichment and significance based on re-sampling permutations within the mappable genome. Results described in this manuscript are based on 100,000 re-sampling permutations.

Accession numbers for comparison datasets are shown in the Key Resources Table. For ChromHMM and ChIP-seq comparison datasets, we downloaded published tables of peak regions from the ENCODE portal or from the Gene Expression Omnibus. Similarly, for analyses of gene sets, we downloaded gene lists from HDSigDB (https://www.hdinhd.org/2018/05/22/hdsigdb/), and we defined the regions of interest as +/- 20 kb of the canonical TSSs of the genes in each set.

### Code and Data Availability

Code used in the analysis of ChIP-seq data is publicly available: https://github.com/seth-ament/ament-carroll-collab. ChIP-seq data have been deposited in the Gene Expression Omnibus (GSE150750). Processed ChIP-seq data, including peak locations (BED files) and genomic coverage (bigWig files), have been deposited in the Neuroscience Multi-Omic Archive (NemO Archive), where we have created a TrackHub, suitable for viewing in the UCSC Genome Browser: http://data.nemoarchive.org/other/grant/sament/sament/htt-chipseq/.

## Acknowledgements

We thank Steven Henikoff, Peter Skene, and Christine Codomo for guidance on X-ChIP-seq protocol optimization, as well as the members of the Carroll lab for discussion regarding the findings and manuscript preparation. This work was supported by the CHDI Foundation (J.B.C., S.A.A.), a Huntington Society of Canada New Pathways grant (J.B.C.), and a National Science Foundation Graduate Research Fellowship to JRP.

## Author Contributions

Conceptualization: JRP, JPC, LEH, NDP, SAA and JBC. Supervision: LEH, NDP, SAA and JBC. Investigation: JPC, DEB, HRP, RMB, SRC. Data Curation, Formal Analysis, and Visualization: ACS, JBC, JPC, and SAA. Funding Acquisition: JBC, SAA. Writing – Original Draft: JBC, JPC, SAA. All authors reviewed and approved the final manuscript.

## Supplementary Information Titles and Legends

**Supplementary Figure 1:**
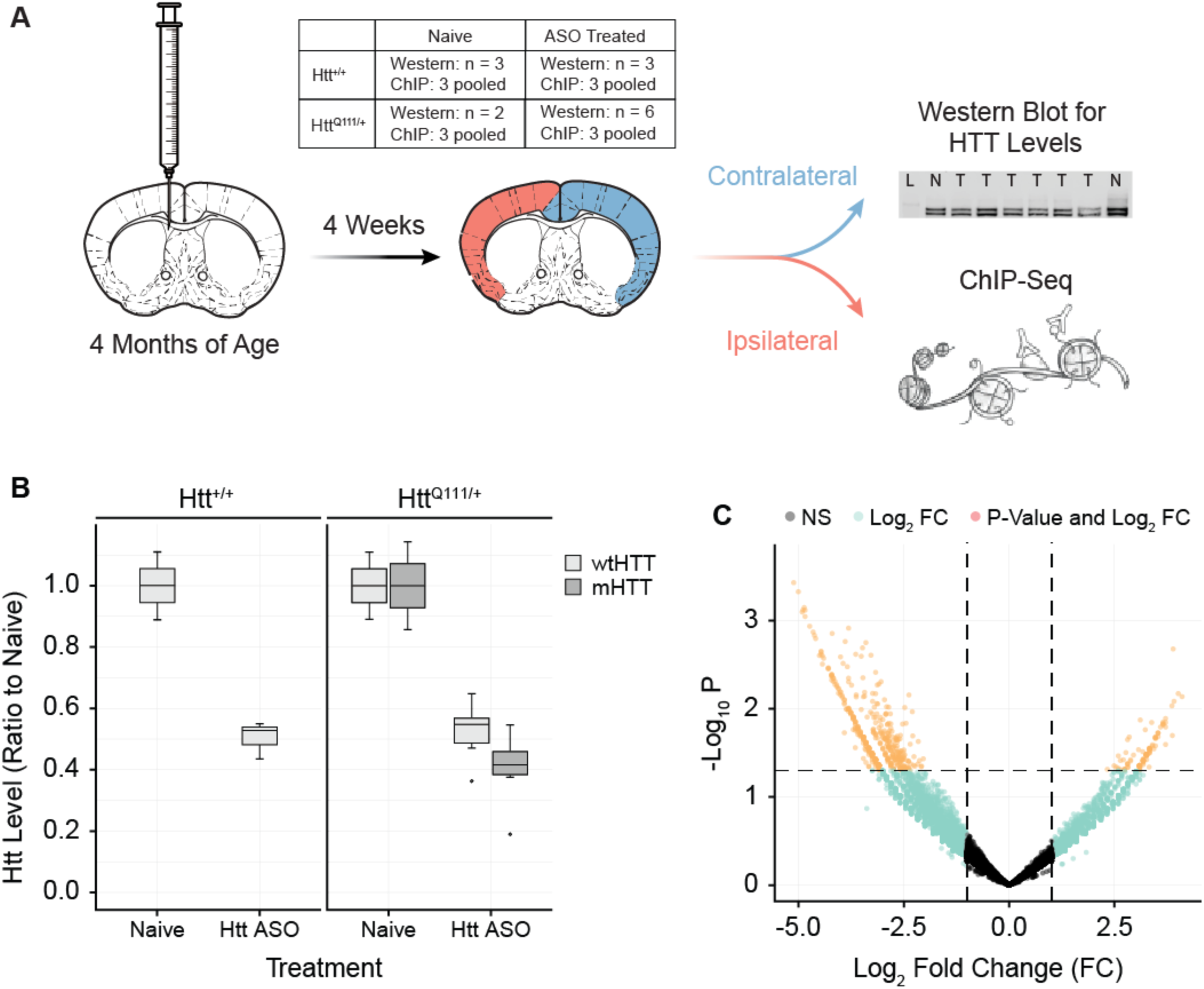
Treatment with a HTT-lowering ASO reduces HTT ChIP-seq peaks. A) Mice underwent unilateral intracerebroventricular injection with HTT-targeted ASO. Cortex ipsi- and contralateral to the injection were collected at four weeks post-injection. HTT lowering was assessed in the contralateral cortex, while ChIP was performed on the ipsilateral cortex. B) Quantification of western blots probed with anti-HTT EPR5526 show HTT lowering of 53% in the contralateral cortex at four weeks post-ASO injection (ANOVA, Treatment effect p = 4.11e09). C) Volcano plot demonstrating 13,820 reproducible HTT-associated peaks in the cortex, 8,308 (60%) showing Log2(fold change) < 0 with ASO treatment. Likelihood that more than 50% had negative fold changes, p = 4.2e126 (binomial test). Of differentially occupied peaks (p < 0.05): 502 out of 622 had negative fold changes (80%) with ASO treatment. Despite the small sample size and partial HTT knockdown, we detected a global depletion of HTT occupancy in the cortex of mice treated with HTT-lowering ASOs.

**Supplementary Table 1 – Significant HTT peak calling results, including proximal gene names, including genotype category labels.** The “MACS Significant HTT Peaks” tab includes the MACS peak calling results - annotated with the most proximal gene and the distance to it. The “Peak.Label” column indicates whether each peak is in the category of “mHTT-specific”, “WT-specific”, or “WT-mHTT-shared” – see text for details on these categories.

**Supplementary Table 2 – HTT Peaks Per Gene.** A per-gene summary of HTT peaks across the categories of “mHTT-specific”, “WT-specific”, or “WT-mHTT-shared.”

**Supplementary Table 3 – HTT DiffBind.** Differential HTT occupancy in *Htt^Q111/Q111^ vs. Htt^+/+^.* Used to generate volcano plot Fig. 2A.

**Supplementary Table 4 – HTT DiffBind Enrichr Results.** Geneset enrichment results for differential HTT peak occupancy in Figs. 2B, 2C.

**Supplementary Table 5 – DiffBind Summary ActiveMotif.** Differential occupancy of EZH2, H3K27ac, H3K27me3, H3K4me3, and H3K9me3, peak regions. These results were used to generate Fig. 3A.

**Supplementary Table 6 – H3K27me3 Enrichment.** Enrichment of H3K27me3 peaks with lower occupancy in *Htt^Q111/+^ vs. Htt^+/+^*striatum (p < 0.005) assessed for enrichment using `Enrichr`. Tabs correspond to terms for Gene Ontology Biological Process (GOBP), Gene Ontology Molecular Function (GOMF), and ENCODE and ChEA Consensus Transcription Factors (ENCODE-CHEA_tfs). Used to generate Figs. 3B-D.

**Supplementary Table 7 – HTT Enrichment ActiveMotif Peaks.** Summary of `GAT` results testing the enrichment of HTT ChIP-seq peaksets amongst EZH2, H3K27ac, H3K27me3, H3K4me3, and H3K9me3, peak regions. These results were used to generate Fig. 3E.

**Supplementary Table 8 - HTT Peaks, ActiveMotif and RNASeq Integration.** Includes integrated ChIP-seq and RNA-seq data for all intervals containing robust HTT peaks. For each HTT peak (“PeakRegion.ID”), available changes in RNA expression of included genes (“Langfelder et. al. RNA-Seq”) and our other ChIP-seq data (“Re-analysis of ActiveMotif ChIP-Seq”) are shown. These results were used to generate Fig. 5A-B and 6A-C.

**Supplementary Table 9 - HTT HDSigDB Overlap.** Includes enrichments for HTT Peak sets and the genes included in each HDSigDB gene set. HDSigDB gene set meta information is included on the tab “HDSigDB.Genesets”. These results were used to generate Fig. 5C.

